# Development of a Freeze-Dried CRISPR-Cas12 Sensor for Detecting *Wolbachia* in the Secondary Science Classroom

**DOI:** 10.1101/2021.10.06.463384

**Authors:** Grant A. Rybnicky, Radeen A. Dixon, Robert M. Kuhn, Ashty S. Karim, Michael C. Jewett

## Abstract

Training the future synthetic biology workforce requires opportunity and exposure to biotechnology concepts and activities in secondary education. Detecting *Wolbachia* bacteria in arthropods using PCR has become a common way for secondary students to investigate and apply DNA technology in the science classroom. Despite this framework, cutting-edge biotechnologies like CRISPR-based diagnostics have yet to be widely implemented in the classroom. To address this gap, we present a freeze-dried CRISPR-Cas12 sensing reaction to complement traditional DNA technology education and teach synthetic biology concepts. The reactions accurately detect *Wolbachia* from arthropod-derived PCR samples in under 2 hours and can be stored at room temperature for over a month without appreciable degradation. The reactions are easy-to-use and cost less than $40 to implement for a classroom of 22 students including the cost of reusable equipment. We see this technology as an accessible way to incorporate synthetic biology education into existing biology curriculum, which will expand biology educational opportunities in science, technology, engineering, and mathematics (STEM) education.

## Introduction

The global market for synthetic biology products is estimated to grow to over $30 billion USD by 2026 from a current $9.5 billion USD in 2021^1^. To support this growth, a workforce of graduating students must be inspired and trained to work in synthetic biology, and its associated science, technology, engineering, and mathematics (STEM) disciplines^2–5^. This requires student exposure to, and formative hands-on experiences with, STEM educational activities.

Many secondary schools in the United States have adopted STEM-related frameworks to teach science and engineering practices. For example, thousands of secondary science classrooms integrate DNA technology into curriculum through activities like the *Wolbachia* Project^6^. The *Wolbachia* Project teaches ecology, biotechnology, and bioinformatics concepts through surveying native arthropods for *Wolbachia* infection. Given the widespread range of these bacteria globally^7^, detection of *Wolbachia* in the secondary education classroom has become an accessible model for learning about DNA biotechnologies. The project includes collection and identification of arthropods from the students’ local environment, bulk DNA extraction from the specimen, Polymerase Chain Reaction (PCR) amplification of a *Wolbachia* specific amplicon, and gel electrophoresis-based detection of the PCR amplicon (Figure 1A). By carrying out these activities, students learn about the existence of standard molecular biotechnology techniques as well as practice them in an engaging inquiry-based way. While curriculum like this teaches foundational techniques important to synthetic biology, translation of the most cutting-edge synthetic biology concepts has been limited.

**Figure 1.**
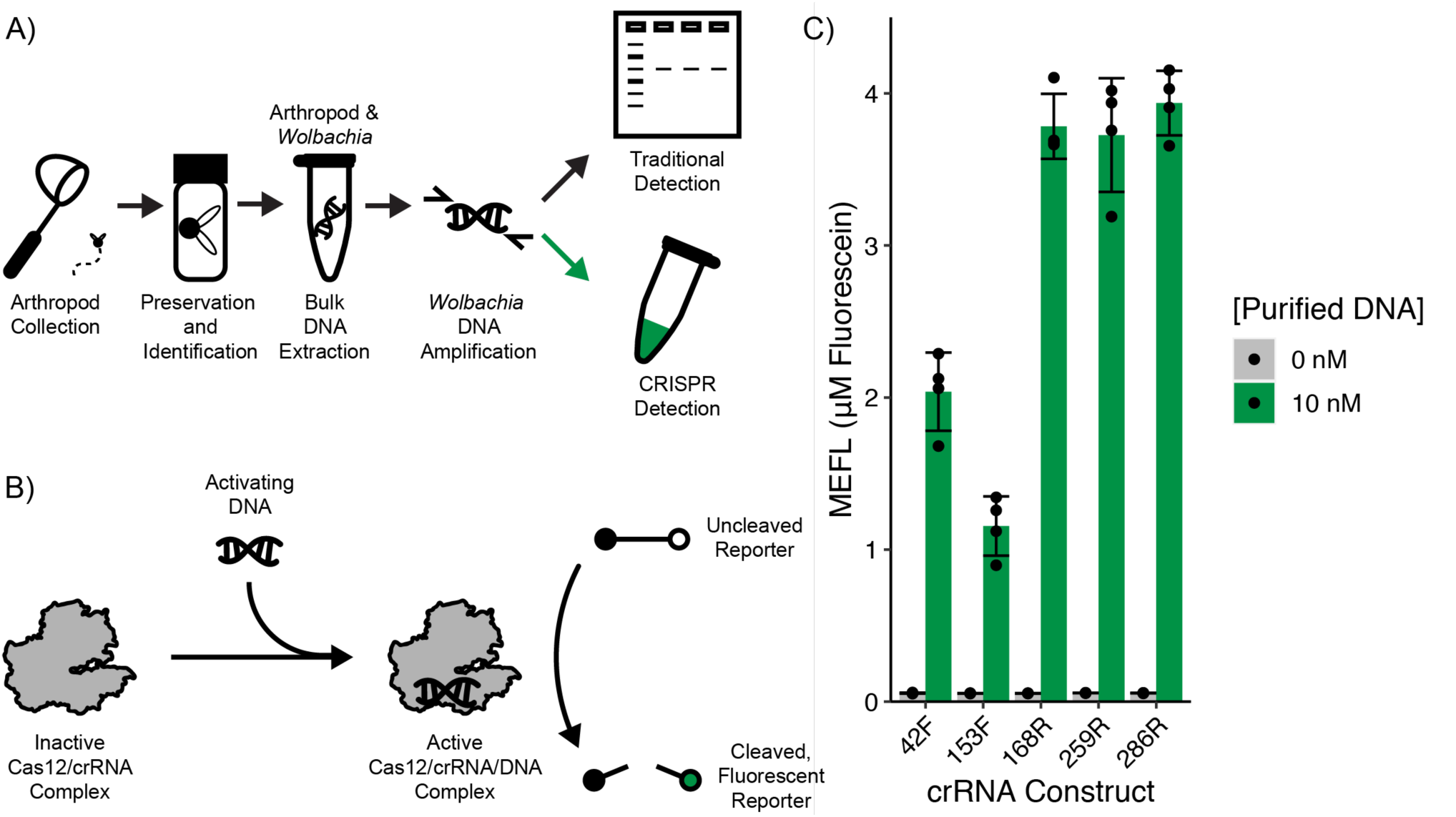
CRISPR-Cas12 diagnostics can be integrated into existing high school biology curriculum. A) Cas12-based DNA sensing reactions can theoretically be used in addition to gel electrophoresis to detect *Wolbachia* infection in student collected arthropod samples. B) Cas12 is activated by the presence of DNA complementary to the crRNA and indiscriminately cleaves ssDNA. A ssDNA quencher-fluorophore probe can convert this activity to observable fluorescence. C) All possible crRNAs against the *Wolbachia* amplicon were tested for activity. Endpoint fluorescence after incubation at 37 °C for 2 hours is reported. Error bars represent standard deviation of 4 replicates. Purified synthetic DNA with the same sequence as the consensus *Wolbachia* amplicon was used as an activator.

A major barrier to teaching synthetic biology and other cutting edge STEM concepts is that education standards can be inflexible and make it difficult to interject new topics not explicit in the standard curriculum^8^. Despite this barrier, recent synthetic biology-based educational efforts like BioBits^2–4^, the BioBuilder Educational Foundation^9^, the International Genetically Engineered Machines competition, Amino Labs, and The ODIN are making progress. Yet, there are opportunities to do more. A particularly exciting area for expansion is in portable cell-free diagnostics^10,11^. In recent years, point-of-use, cell-free diagnostics based on clustered regularly interspaced short palindromic repeats (CRISPR) CRISPR-associated (Cas) technologies have emerged to detect everything from human parasites^12^ to SARS-CoV-2^13,14^. A key feature of CRISPR-based diagnostics is the ability to use Cas12^15^ or Cas 13^16^ to convert DNA or RNA binding into an observable signal, respectively (Figure 1B). Incorporating synthetic biology innovations like this into existing biology lab activities could lead to improved classroom activities but more importantly the integration of synthetic biology education into the educational infrastructure that already exists.

Here, we present a cell-free, tube-based, and freeze-dried CRISPR-Cas12 (FD-CC12) DNA sensing reaction that can be directly integrated into a popular DNA technology educational activity developed by the *Wolbachia* Project to visualize the presence of *Wolbachia* DNA in arthropod-derived samples (Figure 1A). In contrast to gel electrophoresis-based visualization, FD-CC12 reactions do not require specialized electrophoresis equipment, pipetting into small wells, DNA staining or band interpretation, and are ready to interpret in two or fewer hours. FD-CC12 reactions are less technically challenging to perform than gel electrophoresis, and a positive result is easily interpreted as fluorescence visible in the tube after incubation at 37 °C. This system modifies a traditional DNA technology educational activity and incorporates synthetic biology concepts through CRISPR-based diagnostics. In addition, FD-CC12 reactions are cheaper to implement than typical electrophoresis systems, easy to use, and are reliable in detecting *Wolbachia* infection within field-collected arthropod specimens. The reactions do not require refrigeration and are stable at room temperature for over a month. Using this FD-CC12 *Wolbachia* assay, secondary science students will gain hands-on experience with cutting edge synthetic biology and stable, low cost, reliable *Wolbachia* detection in arthropods. We anticipate that integrating synthetic biology into existing activities will be a powerful approach to teaching innovation in synthetic biology.

## Results

### Development of a Cas12 Diagnostic to Detect *Wolbachia*

We set out to develop a CRISPR-Cas12 diagnostic for *Wolbachia* detection to implement in secondary classrooms. First, we developed a functional CRISPR-Cas12 diagnostic. Cas12, like other CRISPR nucleases, binds and cleaves DNA at a specific sequence encoded by a CRISPR RNA (crRNA)^17^. Once activated, Cas12 indiscriminately cleaves single stranded DNA (ssDNA) in solution (Figure 1B)^15^. This indiscriminate cleavage activity can be visualized within a diagnostic in a variety of ways, including the use of a ssDNA chemically modified with a fluorophore on one end and a fluorescence quencher on the other. In the uncleaved state, the ssDNA probe emits no fluorescence upon excitation as the quencher molecule is near the fluorophore and absorbs the fluorescent emission from the fluorophore. Once cleaved, the ssDNA probe is no longer intact, and the fluorescence quencher is not close enough to the fluorophore to absorb the fluorescent emission. To test for CRISPR-Cas12 activity, we screened all possible crRNAs that can target the consensus 483 bp *Wolbachia* amplicon previously developed for the *Wolbachia* Project. Of the five locations with the appropriate TTTV Protospacer Adjacent Motif (PAM)^17^ (Supplementary Figure S1), all possible crRNAs mediated detection of the *Wolbachia* amplicon (Figure 1C). Although all were functional, reaction endpoint fluorescence varied with different crRNA constructs. crRNA 286R yielded the highest endpoint fluorescence as well as fold activation. Thus, we selected crRNA 286R for further development.

### Characterization of Freeze-Dried Cas12 Diagnostics

With a functional CRISPR-Cas12 diagnostic for *Wolbachia* detection in hand, we next wanted to assess the possibility of freeze-drying the system. Previous works have shown that cell-free systems, including CRISPR-based diagnostics, can be freeze-dried for increasing stability and portability^18,19^. Such features would be advantageous for preparation and delivery in the classroom setting. We freeze-dried CRISPR-Cas12 cell-free reactions in a VirTis BenchTop Pro lyophilizer at ≤100 mtorr and −80 °C overnight or until fully freeze-dried. We then compared CRISPR-Cas12 reactions prepared from fresh reagents and these pre-assembled, lyophilized reactions to determine whether the CRISPR-Cas12 formulation is stable through the lyophilization process. Unfortunately, when lyophilized, the CRISPR-Cas12 formulation demonstrated no activity in the presence of activating DNA (Figure 2A). We hypothesized that the lack of activity was due to degradation of the Cas12 protein or the crRNA.

**Figure 2.**
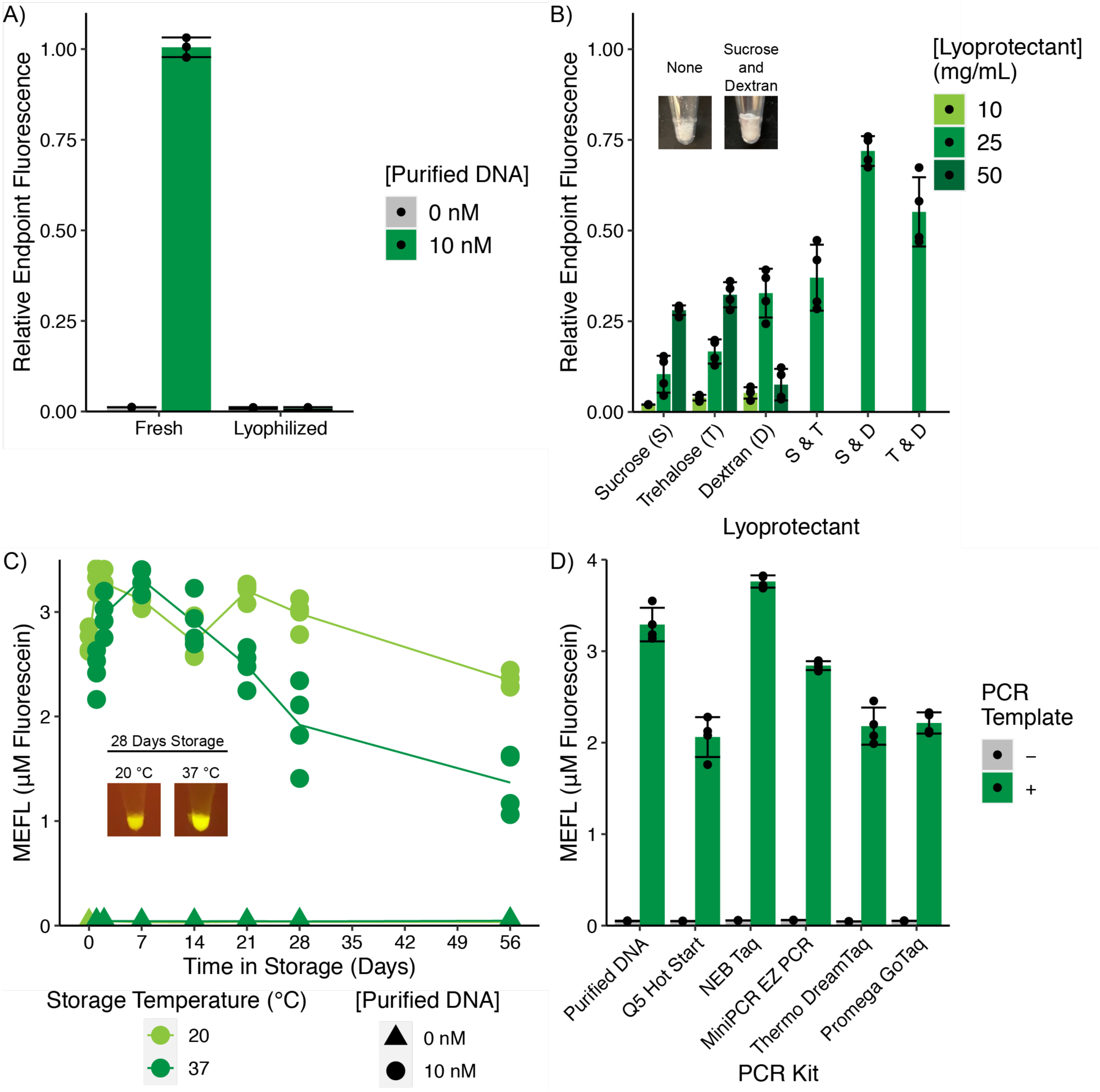
Development and characterization of freeze-dried CRISPR-Cas12 sensing reactions. Endpoint fluorescence was determined after incubation at 37 °C for 2 hours. Error bars represent standard deviation of 4 replicates. B) Inset photos are representative images of FD-CC12 reaction prior to rehydration. C) Lyophilized reactions supplemented with 25 mg/mL each of sucrose and dextran were stored in PCR tubes vacuum sealed in Food Saver® bags with a desiccant card. All data from 4 replicates of each reaction are plotted. The line connects the average value at each time point. Inset photos are representative images of reactions rehydrated with purified DNA and incubated at 37 °C for 2 hours viewed with a portable blue light imager. D) Lyophilized reactions supplemented with 25 mg/mL each of sucrose and dextran were rehydrated with the same volume of purified DNA or unpurified PCR product as the initial reaction volume prior to lyophilization.

To address this bottleneck and improve the stability of the reactions, we evaluated diagnostic activity when reactions were lyophilized in the presence of different lyoprotectant formulations (Figure 2B). Specifically, we tested sucrose, trehalose, and dextran (70 kDa average weight) individually and in combination. We found that the activity of reactions supplemented with sucrose and trehalose individually increased with concentration, but dextran exhibited a concentration optimum of 25 mg/mL in the final reaction. Although combining 25 mg/mL each of sucrose and trehalose achieved similar endpoint fluorescence as 50 mg/mL of each respective lyoprotectant, combining 25 mg/mL of either sugar with 25 mg/mL dextran increased the retained activity. Combining 25 mg/mL each of sucrose and dextran exhibited approximately 75% recovery of the pre-lyophilization signal.

Additionally, all formulations containing dextran exhibited beneficial changes in freeze-dried pellet consistency. Rather than a powdery pellet subject to movement by static electricity, all lyophilized dextran formulations exhibited a bulky, solid pellet unaffected by static charges. This made manipulation of pellets much easier and decreased the risk of static electricity ejecting portions of the pellet from the tube. In making the pellet easier to work with, the inclusion of dextran in the lyoprotectant formulation made the FD-CC12 reactions more amenable to use in secondary science classrooms. Moving forward we chose to use 25 mg/mL each of sucrose and dextran to formulate our freeze-dried reactions.

We next characterized the final formulation on freeze-dried CRISPR-Cas12 (FD-CC12) reaction stability over time. After lyophilization we stored reactions at either 20 or 37 °C in vacuum sealed bags with a desiccant card for two months, monitoring activity over time. When stored at room temperature, the lyophilized reactions retained approximately 75% activity in the on state after 2 months of storage without appreciable leak in off state (Figure 2C). When stored at 37 °C for the same amount of time, the reactions retained over 40% activity in the on state without leak.

### Assessment of Freeze-Dried Cas12 Diagnostics with different PCR mixes

In the *Wolbachia* Project, students amplify DNA using PCR following DNA extraction from arthropods. Different classrooms may use different PCR reaction mixtures, based on price and preference. To assess generalizability of our FD-CC12 reactions and facilitate integration into existing classroom activities, we therefore tested compatibility with common PCR reaction mixtures. We evaluated whether several commonly used PCR kits (Q5 Hotstart, NEB Taq, MiniPCR EZ PCR, Thermo DreamTaq, and Promega GoTaq) could be used to rehydrate the FD-CC12 reactions and maintain functionality. Final endpoint fluorescence varied among FD-CC12 reactions rehydrated with different PCR kits, all achieving at least 60% of the signal achieved by reactions rehydrated with purified DNA (Figure 2D). Despite this variability in endpoint fluorescence, all the reactions produced observable signal in the on state and negligible signal in the off state. These data highlight the robustness of our FD-CC12 diagnostic for detecting *Wolbachia*.

### Assessment of Freeze-Dried Cas12 Diagnostics in the Classroom

With stable and robust FD-CC12 reactions established, we next wanted to test compatibility with the *Wolbachia* Project educational framework. We first wanted to make sure that *Wolbachia* DNA from arthropod-derived samples could be detected. When rehydrated with a PCR reaction that used bulk DNA from a *Wolbachia* infected *Nasonia vitripennis* wasp as template, the FD-CC12 reaction achieved endpoint yields comparable to those observed in the PCR kit screen (Figure 3A). Using a *Wolbachia*-free *N. vitripennis* wasp-derived PCR, the FD-CC12 reaction showed signal above background observable by fluorescent plate reader but not by eye.

**Figure 3.**
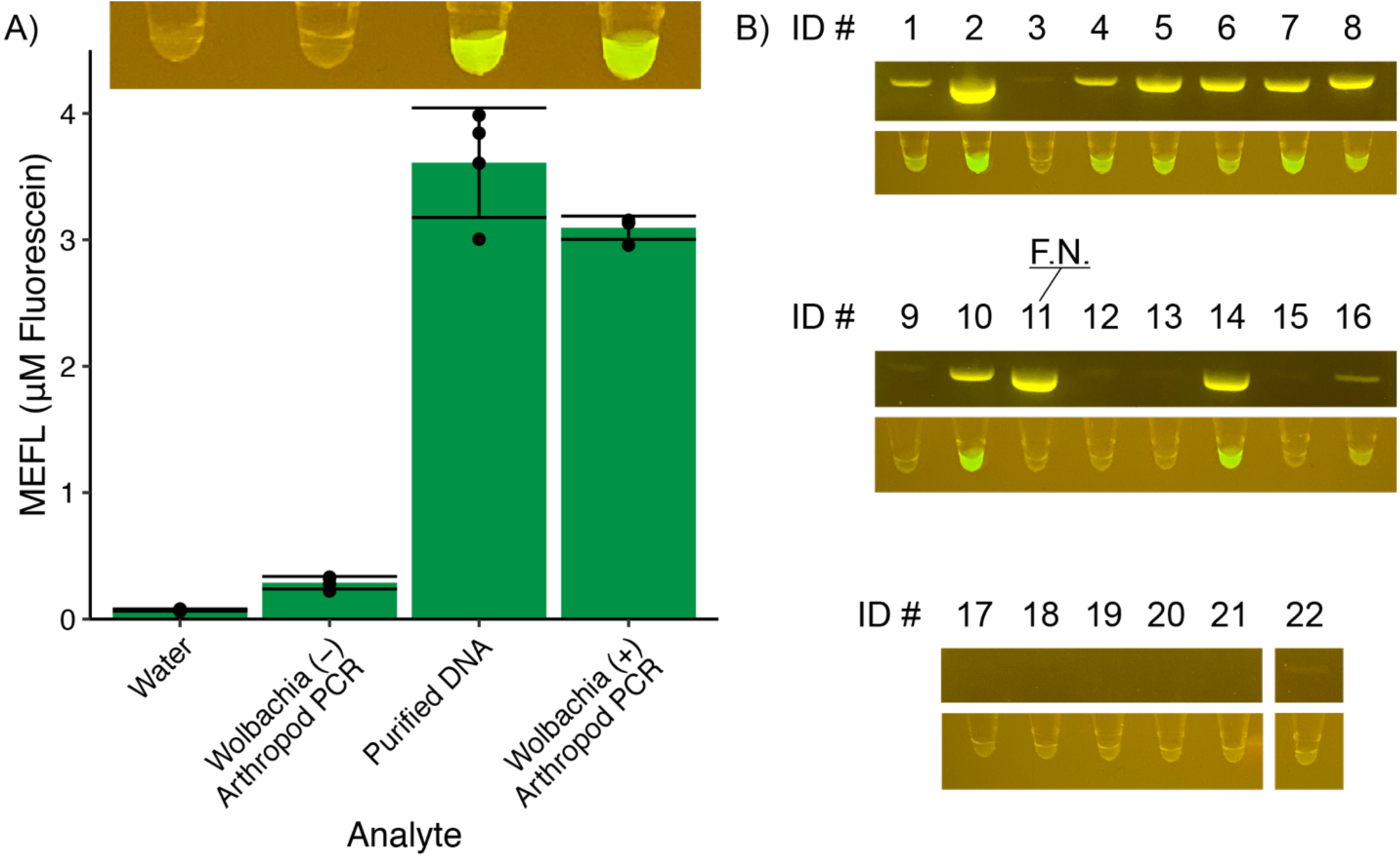
Freeze-dried CRISPR-Cas12 reactions can be used by a high school student to detect *Wolbachia* infection in field-collected arthropods. A) The inset photo is representative of the control reactions incubated at 37 °C for 2 hours viewed with a portable blue light imager. B) Arthropods were collected from two locations, Brevard, NC & Roswell, GA. A high school student extracted DNA from the specimens, amplified *Wolbachia* DNA using PCR, and concurrently used gel electrophoresis and FD-CC12 reactions to detect the presence of the *Wolbachia* amplicon. Both the gel and Cas12 reactions were imaged using a portable blue light imager.

Once validated with control insects, we next assessed the ability of the workflow to be completed in a high school classroom setting by a high school student. FD-CC12 reactions were prepared at Northwestern University and shipped to Centennial High School (CHS) in Roswell, Georgia. At CHS, a student collected and identified 22 arthropods across 6 orders (Supplementary Table S2), extracted bulk DNA from each specimen, and performed PCR to amplify the *Wolbachia* amplicon as is standard practice in the *Wolbachia* Project workflow. Those PCR products were then used to rehydrate FD-CC12 reactions and were also visualized using gel electrophoresis. Out of the 22 samples tested, 21 agreed between FD-CC12 reaction and gel electrophoresis (Figure 3B). For sample ID# 11, the gel indicated *Wolbachia* infection while the FD-CC12 reaction indicated no infection. Upon sequencing the *Wolbachia* amplicon for sample ID# 11, it was revealed that there were 5 mutations within the protospacer region the crRNA 286R targeted (Supplementary Figure S2). As *Wolbachia* are a diverse genus of bacteria, it is not unreasonable to expect that some isolates would evade detection by any single crRNA. Using multiple crRNAs in the same reaction to create OR gate logic, it may be possible to detect a wider diversity of isolates. Given the compatibility of the FD-CC12 reactions with arthropod-derived samples, successful completion by a high school student, and high accuracy of sensing reactions, this demonstrates that FD-CC12 reactions can be integrated into the *Wolbachia* Project laboratory activities.

## Discussion

We developed a FD-CC12 *Wolbachia* DNA sensing reaction that can be used to teach synthetic biology concepts in the secondary science classroom. As designed, the sensing reactions detect a PCR amplicon commonly used to diagnose *Wolbachia* infection in field-collected arthropods as a demonstration of DNA technologies (Figure 1A). We identified and evaluated 5 functional crRNAs that can be used to detect this amplicon with Cas12 (Figure 1C). Additionally, we identified a lyoprotectant additive formulation that both recovers about 75% of reaction endpoint fluorescence after lyophilization (Figure 2A&B) and is stable at room temperature for over a month without appreciable degradation (Figure 2C). Using this formulation, we evaluated the reaction’s compatibility with commonly used PCR kits, finding that the freeze-dried reactions could be rehydrated with virtually any unpurified PCR product and still distinguish the presence of the *Wolbachia* amplicon in the sample (Figure 2D). To validate the reaction can be applied to the established classroom activity, we rehydrated freeze-dried reactions with arthropod-derived samples. *Nasonia* wasps with known *Wolbachia* infection states yielded results as expected (Figure 3A). Screening of 22 field-collected arthropods across 6 orders for *Wolbachia* infection by a high school student using gel electrophoresis and FD-CC12 reactions yielded a 95% accuracy rate (Figure 3B).

While we recommend that FD-CC12 sensing reactions be used to complement gel electrophoresis in DNA technology education, we do recognize that specific features of our sensing reactions may make them a more attractive readout than gel electrophoresis to secondary science educators. The FD-CC12 reactions primarily lower two barriers DNA visualization in the classroom: 1) FD-CC12 reactions are less technically difficult to set up and run than gel electrophoresis and 2) FD-CC12 reactions are less expensive. While commonplace in the molecular biology research lab, gel electrophoresis poses a few technical hurdles to implementation that science educators may face. Foremost, gel electrophoresis is intimidating to educators who have not had hands-on training in a laboratory or elsewhere. The ability to properly prepare and cast an agarose gel, pipette samples into the well of a gel, and consistently stain and visualize gels takes hands-on practice before it becomes routine. Collectively, these technical hurdles may cause teachers to seek more expensive alternatives (precast gels for example) or to avoid implementation of gel electrophoresis altogether. Alternatively, the FD-CC12 reactions contain all necessary components within a single tube, are easier to pipette analyte into, and are visualized by handheld blue light imager without the addition of stains. The use of freeze-dried cell-free reactions has the potential to increase teacher confidence in their laboratory skills and encourage them to attempt laboratory activities they may have been hesitant to try before like gel electrophoresis. Similarly, the initial monetary investment required to implement FD-CC12 sensing in the classroom is lower than implementing gel electrophoresis. Requiring only incubation at 37 °C and a blue light imager like the MiniPCR P51™ Molecular Fluorescence Viewer, the FD-CC12 reactions have a startup cost of ~$38 for a 22-student class. Alternatively, the equipment to cast gels, generate current, house the gel in buffer, and visualize the gel as well as initial reagents cost ~$1000 or ~$300 for the same sized class for traditional or education optimized gel electrophoresis equipment, respectively (Table 1). Assuming one sample per student and class size of 22 students, the CRISPR-Cas12-based activity is cheaper to run for 848 students or fewer than education optimized gel electrophoresis set up. The price of FD-CC12 reactions can be further reduced about 10X by applying state-of-the-art cell-free technology like the use of paper instead of tubes^20^. That being said, gel electrophoresis systems are generalizable to visualizing almost any DNA sample while the FD-CC12 sensing reaction presented is specific to *Wolbachia* amplicon detection given the crRNA used.

**Table 1.** Comparison of DNA visualization method equipment and consumable costs. Cost calculation assumes 22 samples and 2 DNA ladders per gel. Comparison does not include the cost of an incubator to incubate rehydrate FD-CC12 reactions as the PCR machine used to generate the PCR product can be used to heat the reaction.

| Setup | Component | Supplier | Product Number | Unit Cost | Initial Equipment Investment | Reagent Cost/ Reaction |
| --- | --- | --- | --- | --- | --- | --- |
| Traditional Gel Electrophoresis | Casting Tray | BioRad | 1640300 | \$993 | \$993.00 | \$0.31 |
|  | Electrophoresis Chamber |  |  |  |  |  |
|  | Power Supply |  |  |  |  |  |
| | Agarose | BioRad | 1613100 | \$0.84 / gel | | |
| | TBE Buffer | BioRad | 1610770 | \$0.66 / gel | | |
| | FlashBlue Stain | EDVOTEK | 609 | \$2.63 / gel | | |
| | DNA Ladder | BioRad | 1708352 | \$2.76 / gel | | |
| Education Optimized Gel Electrophoresis | blueGel™ electrophoresis with built-in transilluminator | MiniPCR | QP-1500-01 | \$300.00 | \$300.00 | \$0.15 |
| | GelGreen® Agarose Tabs™ | MiniPCR | RG-1500-10 | \$1.20 / gel | | |
| | TBE Buffer | MiniPCR | RG-1502-02 | \$0.09 / gel | | |
| | DNA Ladder | MiniPCR | RG-1003-01 | \$2.00 / gel | | |
| 10 µL Tube-Based FD-CC12 | P51™ Molecular Fluorescence Viewer | MiniPCR | QP-1900-01 | \$28.00 | \$28.00 | \$0.47 |
| | Freeze-Dried Reaction | See Table S3 | See Table S3 | \$0.2907 / reaction | | |
| | PCR Tube | Thermo Scientific | AB2000 | \$0.18 / reaction | | |
| 1.8 µL Paper-Based FD-CC12 | P51™ Molecular Fluorescence Viewer | MiniPCR | QP-1900-01 | \$28.00 | \$28.00 | \$0.05 |
| | Freeze-Dried Reaction | See Table S3 | See Table S3 | \$0.0523 / reaction | | |
| | Filter Paper | Cole-Parmer | UX-06647-20 | \$0.0007 / reaction | | |

Although we focused on detection of *Wolbachia* in the secondary science classroom, we note that the technology described has potential applications beyond the scope presented. Within science education, the DNA that is detected can easily be changed by altering the crRNA in the reaction. There are numerous well established laboratory activities that use gel electrophoresis to detect double stranded DNA PCR product. Given the presence of a functional PAM sequence in the DNA analyte and appropriate design of a crRNA, the technology described here can be extended to other educational activities beyond *Wolbachia* detection. The technology is not limited to the classroom either. As the freeze-dried cell-free reactions are still functional after 56 days storage at 37 °C (Figure 2C), it is feasible to use these reactions in on-site biological field research. In addition, FD-CC12 reactions could be implemented in field stations where gel electrophoresis may be difficult or impractical. As mentioned before, other DNAs can alternatively be detected by exchanging the crRNA in the system. This can lead to applications such as tracking of specific organisms in the environment by DNA detection, identification of visibly indistinguishable species, or monitoring the spread of mobile genetic elements in the environment.

Taken together, FD-CC12 reactions have the potential to be a simple, cost-effective companion to gel electrophoresis that provide a hands-on way to teach CRISPR-based nucleic acid detection and concepts of synthetic biology. In integrating this activity with the established DNA technology workflow designed by the *Wolbachia* Project, we anticipate widespread adoption of CRISPR-Cas12-based detection into the classroom, further strengthening the effort to expose school age children to synthetic biology concepts and priming the future STEM workforce.

## Methods

### Cas12 Sensing Reactions

Fresh Cas12 reactions were assembled by mixing reaction buffer (40 mM Tris-HCl, 60 mM NaCl, 6 mM MgCl2, pH 7.3), 45 nM AsCas12a V3 (IDT), 45 nM crRNA (IDT), 1 µM ssDNA Fluorophore-Quencher (FQ) Probe (IDT), and lyoprotectant. Prior to addition of FQ Probe and lyoprotectant, the mixture was equilibrated at 37 °C for 15 minutes. Upon addition of the analyte, the reaction was incubated at 37 °C and fluorescence was measured (490 nm Excitation, 525 nm Emission) using a Synergy H1 microplate reader (BioTek, USA) and Gen5 v. 2.09 (BioTek) software.

Lyophilized Cas12 reactions were assembled as described above and then flash frozen with liquid nitrogen in PCR strip tubes (Thermo Scientific, AB2000) with a small hole melted in the cap of each tube. The reactions were lyophilized using a VirTis BenchTop Pro lyophilizer (SP Scientific) at ≤100 mtorr and −80 °C overnight or until fully freeze-dried. Following lyophilization, tubes were packaged in Food Saver® bags with a desiccant card, vacuum sealed and stored at the indicated temperature. When ready for use, tubes were removed from the Food Saver® bags and rehydrated with the same volume of analyte as the mixture prior to lyophilization. Reactions were incubated at 37 °C and fluorescence was measured as described above.

### Arthropod Collection and *Wolbachia* DNA Amplification

Arthropods belonging to six different orders (Supplemental Table S2) were collected from Roswell, GA and Brevard, NC. Each specimen was preserved in 75% EtOH in a 1.5mL collection tube and frozen at −20 °C. Samples were visually identified to taxonomic order and later identified further by COI barcoding. PCR using the primers described by Folmer^21^ was performed and the product was sanger sequenced (GeneWiz). Consensus sequences for each specimen were obtained using DNA Subway (CyVerse/DNA learning Center) and NIH nucleotide BLAST^22^ was used to taxonomically identify the arthropod. *Wolbachia*-infected and *Wolbachia*-free *Nasonia vitripennis* wasps were obtained from the *Wolbachia* Project through the Bordenstein Lab at Vanderbilt University. Bulk DNA was extracted from each specimen by manual grinding using sterile, plastic minipestles for 2 minutes in a 1.5 mL microcentrifuge tube containing 200 µL of lysis buffer/EDTA mix and subsequent purification with the Monarch Genomic DNA Purification Kit (catalog #T3010) as directed. A 100µL DNA elution was collected stored at −20 °C until used for polymerase chain reaction (PCR).

PCR was performed using 2µL of eluted DNA, Wspec-F (5’−CAT ACC TAT TCG AAG GGA TAG−3’) and Wspec-R (5’−AGC TTC GAG TGA AAC CAA TTC−3’) primers and MiniPCR EZ PCR Master Mix (catalog RG-1000-01). MiniPCR Mini8 thermocyclers were used for PCR with the following protocol: initial denaturation at 94 °C for 120 sec and 34 cycles of denaturation 95 °C for 30 sec, annealing 55 °C for 45 sec, extension 72 °C for 60 sec and final extension 72 °C for 600 sec. PCR product was frozen at −20 °C for further use.

### Gel Electrophoresis and Cas12 Detection of *Wolbachia* DNA in the Classroom

Gel electrophoresis assays were completed in a secondary science education classroom at Centennial High School in Roswell, GA using the MiniPCR blueGel™ electrophoresis system with built-in transilluminator. Gels were poured using GelGreen® Agarose Tabs™ to a concentration of 2% agarose. 5 µL of each PCR product was loaded into a well and were electrophoresed for 30 min. The presence of *Wolbachia* in the arthropod specimen was determined by presence of a 438 bp amplicon using the built in blue light transilluminator.

Freeze-Dried Cas12 reactions were assembled and packaged at Northwestern University in Evanston, IL. The reactions were then shipped at room temperature to Centennial High School in Roswell, GA. Reactions were stored at room temperature for 9 days prior to rehydration. PCR strip tubes containing a freeze-dried pellet of the CRISPR-Cas12 system were rehydrated using 10 µL of PCR product. Tubes were spun briefly at 10,000 rpm and moved to a heat block to incubate at 37 °C for 2 hours. Fluorescence was detected using a MiniPCR p51 fluorescence viewer and photographs taken using a blue light transilluminator.

## Abbreviations

PCR: Polymerase Chain Reaction
CRISPR: Clustered Regularly Interspaced Short Palindromic Repeats
Cas: CRISPR Associated Protein
FD-CC12: Freeze-Dried CRISPR-Cas12
STEM: Science Technology Engineering and Mathematics
K-12: Kindergarten through Twelfth Grade
ssDNA: Single Stranded DNA
FQ: Fluorophore Quencher
PAM: Protospacer Adjacent Motif cr
crRNA: CRISPR RNA

## Author Information

Grant A. Rybnicky - Interdisciplinary Biological Sciences Graduate Program, Northwestern University, 2205 Tech Drive, Hogan Hall 2100, Evanston, Illinois 60208, United States; Chemistry of Life Processes Institute, Northwestern University, 2170 Campus Drive, Evanston, Illinois 60208-3120, United States; Center for Synthetic Biology, Northwestern University, 2145 Sheridan Road, Technological Institute E136, Evanston, Illinois 60208-3120, United States

Radeen A. Dixon - Student, Centennial High School, 9310 Scott Rd, Roswell, GA 30076, United States

Robert M. Kuhn - Teacher, Centennial High School, 9310 Scott Rd, Roswell, GA 30076, United States; Teacher, Innovation Academy Fulton County Schools STEM Magnet High School, 125 Milton Ave. Alpharetta, GA 30009, United States

Ashty S. Karim - Department of Chemical and Biological Engineering, Northwestern University, 2145 Sheridan Road, Technological Institute E136, Evanston, Illinois 60208-3120, United States; Chemistry of Life Processes Institute, Northwestern University, 2170 Campus Drive, Evanston, Illinois 60208-3120, United States; Center for Synthetic Biology, Northwestern University, 2145 Sheridan Road, Technological Institute E136, Evanston, Illinois 60208-3120, United States

Michael C. Jewett - Department of Chemical and Biological Engineering, Northwestern University, 2145 Sheridan Road, Technological Institute E136, Evanston, Illinois 60208-3120, United States; Chemistry of Life Processes Institute, Northwestern University, 2170 Campus Drive, Evanston, Illinois 60208-3120, United States; Center for Synthetic Biology, Northwestern University, 2145 Sheridan Road, Technological Institute E136, Evanston, Illinois 60208-3120, United States; Member, Robert H. Lurie Comprehensive Cancer Center, Northwestern University, 676 N. St. Clair Street, Suite 1200, Chicago, Illinois 60611-3068, United States; Simpson Querrey Institute, Northwestern University, 303 E. Superior Street, Suite 11-131, Chicago, Illinois 60611-2875, United States

## Author Contribution

GAR, RAD, RMK, and MCJ conceived the work presented. GAR, RAD, and RMK ran experiments. GAR, RAD, RMK, ASK and MCJ wrote the paper.

## Acknowledgement

We would like to acknowledge MiniPCR for their contribution of materials and reagents. We would also like to thank Sarah Bordenstein and the Wolbachia Project for helpful conversation and for *N. vitripennis* specimens. We would also like to acknowledge the Centennial High School DNA Club for providing collaborative biology research opportunities for high school students. The authors would like to acknowledge members of the Jewett Lab for helpful discussions. M.C.J. acknowledges support from the Department of Energy Grants DE-SC0018249, DE-NA0003525, and 8J-30009-0029A. the DOE Joint Genome Institute ETOP program, the Office of Energy Efficiency and Renewable Energy Grant DE-EE0008343, the David and Lucile Packard Foundation, the Camille Dreyfus Teacher-Scholar Program, the Defense Threat Reduction Agency Grants HDTRA1-15-10052/P00001 and HDTRA-12-01-0004, the Army Research Office Grants W911NF-20-1-0195, W911NF-18-1-0200, and W911NF-16-1-0372, the National Science Foundation Grants 1936789 and 1844336, and the Air Force Office for Research Grant FA2386-21-1-4078, Army Contracting Command Contract W52P1J-21-9-3023. The work conducted by the U.S. Department of Energy Joint Genome Institute, a DOE Office of Science User Facility, is supported by the Office of Science of the U.S. Department of Energy under Contract No. DE-AC02-05CH11231. G.A.R was supported by the National Science Foundation Graduate Research Fellowship Program under Grant No. DGE-1842165.

