## Supplemental Tables and Figures for "Development of a Freeze-Dried CRISPR-Cas12 Sensor for Detecting *Wolbachia* in the Secondary Science Classroom"

**Supplementary Table S1. List of all nucleic acids used in this study.**

| Name | Nucleic Acid | Sequence (5' to 3') | Origin Reference |
|---|---|---|---|
| ssDNA FQ-probe | ssDNA | /56-FAM/TTATT/3IABkFQ/ | [1] |
| *Wolbachia* Amplicon | dsDNA | https://benchling.com/s/seq-4z2hwvLIoJRraq8vrYRp | [2] |
| Wspec-F | ssDNA | CATACCTATTCGAAGGGATAGGGTCGGTTCGGC | [2] |
| Wspec-R | ssDNA | AGCTTCGAGTGAAACCAATTC | [2] |
| Folmer-F | ssDNA | TGTAAAACGACGGCCAGTGGTCAACAAATCATAAAGATATTGG | [3] |
| Folmer-R | ssDNA | CAGGAAACAGCTATGACTAAACTTCAGGGTGACCAAAAAATCA | [3] |
| crRNA 42F | ssRNA | /AltR1/rUrArArUrUrUrCrUrArCrUrCrUrUrGrUrArGrArUrArCrArCrArGrGrUrGrUrUrGrCrArUrGrGrCrUrGrUrC/AltR2/ | This Study |
| crRNA 153F | ssRNA | /AltR1/rUrArArUrUrUrCrUrArCrUrCrUrUrGrUrArGrArUrArGrGrArGrArCrUrGrCrCrArGrUrGrArUrArArArCrU/AltR2/ | This Study |
| crRNA 169R | ssRNA | /AltR1/rUrArArUrUrUrCrUrArCrUrCrUrUrGrUrArGrArUrUrCrArCrUrGrGrCrArGrUrCrUrCrCrUrUrArArArGrU/AltR2/ | This Study |
| crRNA 260R | ssRNA | /AltR1/rUrArArUrUrUrCrUrArCrUrCrUrUrGrUrArGrArUrCrArGrCrCrCrArUrUrGrUrArGrCrCrArCrCrArUrUrG/AltR2/ | This Study |
| crRNA 287R | ssRNA | /AltR1/rUrArArUrUrUrCrUrArCrUrCrUrUrGrUrArGrArUrArGrGrGrArUrUrGrGrCrUrUrArGrCrCrUrCrGrCrGrA/AltR2/ | This Study |

**Supplementary Table S2. Summary of field-collected arthropod specimens.**

| ID # | Collection Location | Visual Identification | COI Identification | Gel Electrophoresis Result | FD-CC12 Result |
|---|---|---|---|---|---|
| 1 | Annapolis, CA | Termite | *Zootermopsis nevadensis* | Positive | Positive |
| 2 | Roswell, GA | Moth | *Eupithecia miserulata* | Positive | Positive |
| 3 | Roswell, GA | Spider | *Agelenopsis pennsylvanica* | Negative | Negative |
| 4 | Brevard, NC | Ant | *Formica subaenescens* | Positive | Positive |
| 5 | Brevard, NC | Ant | *Formica subaenescens* | Positive | Positive |
| 6 | Brevard, NC | Ant | *Formica subaenescens* | Positive | Positive |
| 7 | Atlanta, GA | Moth | *Hypagyrtis unipunctata* | Positive | Positive |
| 8 | Brevard, NC | Ant | *Formica subaenescens* | Positive | Positive |
| 9 | Roswell, GA | Midge | *Chironomus sp* | Negative | Negative |
| 10 | Roswell, GA | Mosquito | *Aedes albopictus* | Positive | Positive |
| 11 | Roswell, GA | Beetle | *Tribolium confusum* | Positive | Negative |
| 12 | Roswell, GA | Moth | *Tetanolita floridana* | Negative | Negative |
| 13 | Roswell, GA | Beetle | *Harpalus pensylvanicus* | Negative | Negative |
| 14 | Brevard, NC | Ant | *Formica subaenescens* | Positive | Positive |
| 15 | Roswell, GA | Midge | *Chironomus sp* | Negative | Negative |
| 16 | Roswell, GA | Moth | *Clepsis peritana* | Positive | Positive |
| 17 | Roswell, GA | Midge | *Chironomus sp* | Negative | Negative |
| 18 | Roswell, GA | Midge | *Chironomus sp* | Negative | Negative |
| 19 | Roswell, GA | Spider | *Phalangiidae sp* | Negative | Negative |
| 20 | Roswell, GA | Midge | *Chironomus sp* | Negative | Negative |
| 21 | Roswell, GA | Midge | Chironomus sp | Negative | Negative |
| 22 | Roswell, GA | Midge | Chironomus sp | Negative | Negative |

**Supplementary Table S3. Cost of Cas12 Wolbachia Sensing Reaction Components.** Components are priced at size scales consistent with what a standard molecular biology lab may purchase and at publicly advertised prices available online. Price per reaction may be decreased with buying components in bulk or with supplier agreements.

| Component | Cost ($)/ 10 µL Reaction | Supplier | Product Number |
|---|---|---|---|
| AsCas12a V3 | 0.0978260870 | IDT | 1081069 |
| crRNA | 0.0213750000 | IDT | See Table S1 |
| FQ Probe | 0.1709000000 | IDT | See Table S1 |
| MgCl2 | 0.0000126710 | Sigma-Aldrich | 63068-1KG |
| NaCl | 0.0000014587 | Sigma-Aldrich | S3014-5KG |
| Tris HCl | 0.0000076116 | Sigma-Aldrich | T5941-100G |
| Sucrose | 0.0000105000 | Sigma-Aldrich | S0389-5KG |
| Dextran 70 | 0.0006000000 | TCI | D1449 |
| **Total** | **0.290733328** | | |

**Supplementary Figure S1. Location of crRNAs within the *Wolbachia* amplicon.** crRNA targets are indicated with green elongated pentagons with text labels. All crRNA targets are downstream of a TTTV PAM. The strand that the crRNA targets is indicated by the direction of the annotation. Image generated with Benchling.

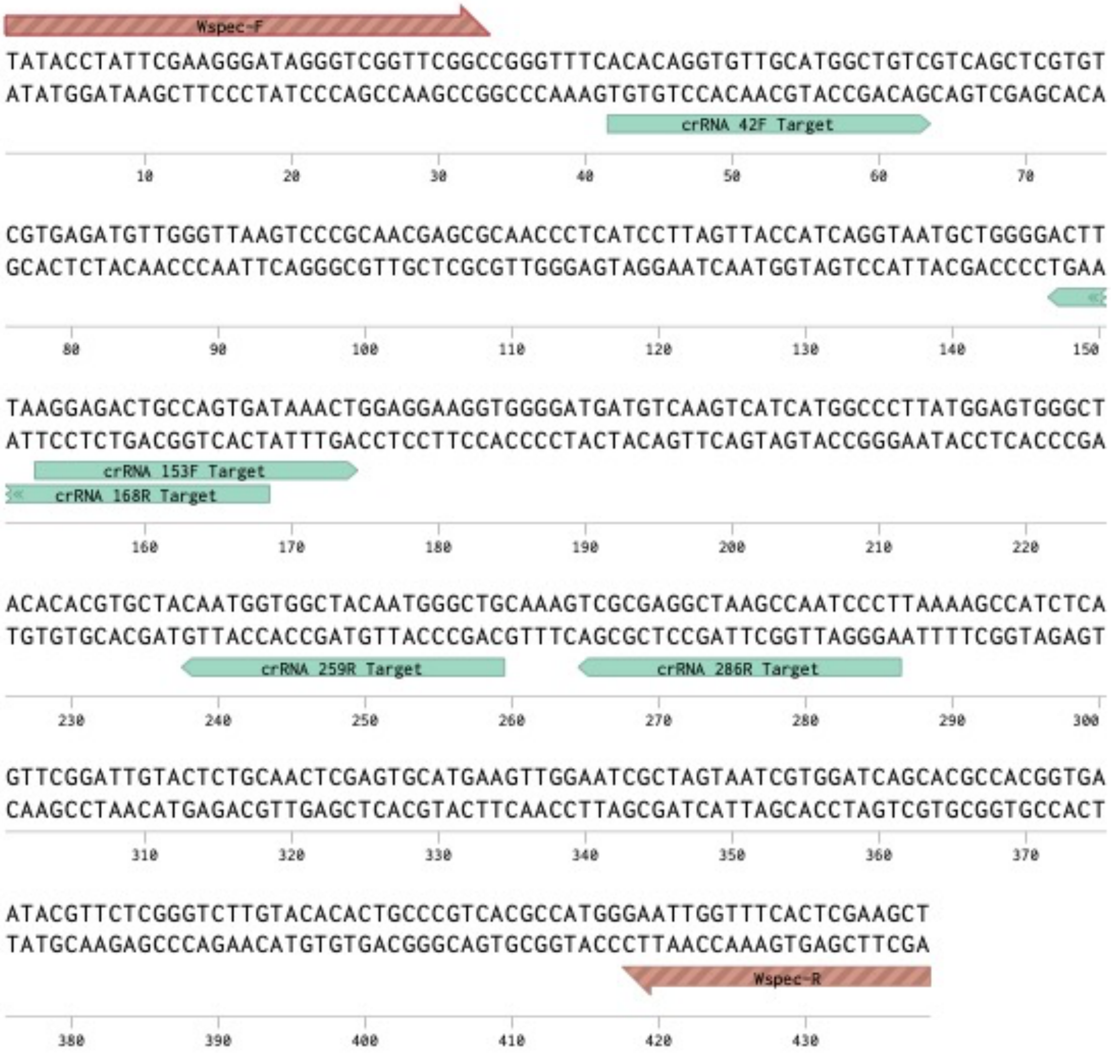

**Supplementary Figure 2. Sequence alignment of consensus *Wolbachia* amplicon and arthropod ID# 11 PCR amplicon.** An excerpt of the consensus *Wolbachia* amplicon sequence is displayed on top and is annotated with the location of the crRNA 286R target. Two sanger sequencing traces and consensus sequences from different sequencing primers are displayed below. Image generated with Benchling.

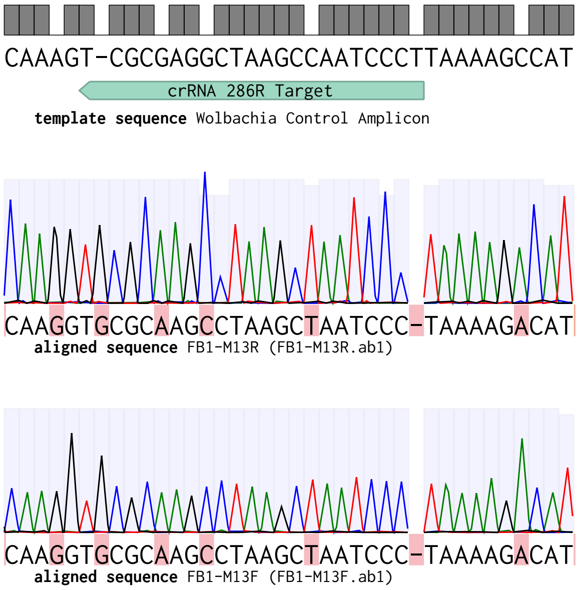

Supplemental Figures References:


(1) Chen, J. S.; Ma, E.; Harrington, L. B.; Da Costa, M.; Tian, X.; Palefsky, J. M.; Doudna, J. A. CRISPR-Cas12a Target Binding Unleashes Indiscriminate Single-Stranded DNase Activity. *Science* **2018**, *360* (6387), 436–439. https://doi.org/10.1126/science.aar6245.

(2) Werren, J. H.; Windsor, D. M. Wolbachia Infection Frequencies in Insects: Evidence of a Global Equilibrium? *Proceedings. Biol. Sci.* **2000**, *267* (1450), 1277–1285. https://doi.org/10.1098/rspb.2000.1139.

(3) Folmer, O.; Black, M.; Hoeh, W.; Lutz, R.; Vrijenhoek, R. DNA Primers for Amplification of Mitochondrial Cytochrome c Oxidase Subunit I from Diverse Metazoan Invertebrates. *Mol. Mar. Biol. Biotechnol.* **1994**, *3* (5), 294–299.
